# Non-contact respiratory measurements of outdoor-housed rhesus macaques (*Macaca mulatta*) using millimeter-wave radar systems

**DOI:** 10.1101/2023.10.11.561971

**Authors:** Toshiki Minami, Daisuke Sanematsu, Itsuki Iwata, Takuya Sakamoto, Masako Myowa

## Abstract

Respiration is an invaluable signal that facilitates the real-time observation of physiological dynamics. In recent years, the advancement of non-contact measurement technology has gained momentum in capturing physiological dynamics in natural settings. This technology is anticipated to find utility in healthcare, not only in humans but also in captive animals, to enhance animal welfare. Currently, the predominant non-contact approach for captive animals involves measuring vital signs through subtle variations in skin color. However, this approach is limited when dealing with body regions covered with hair or outdoor environments under fluctuating sunlight. In contrast, millimeter-wave radar systems, which employ millimeter waves that can penetrate animal fur, exhibit minimal susceptibility to sunlight interference. Thus, this method holds promise for non-contact vital measurements in natural and outdoor settings. In this study, we validated a millimeter-wave radar methodology for capturing respiration in outdoor-housed rhesus macaques (*Macaca mulatta*). The radar was positioned beyond the captive enclosure and maintained at a distance of > 5 m from the target. Millimeter waves were transmitted to the target, and the reflected waves were used to estimate skin surface displacement associated with respiration. The results revealed periodic skin surface displacement, and the estimated respiratory rate was within the reported range of respiratory rates for rhesus macaques. This result suggests the potential applicability of millimeter-wave radar for non-contact respiration monitoring in outdoor-housed macaques. The continued advancement of non-contact vital measurement technology will contribute to the mental and physical monitoring of captive animals to establish comfortable captive environments.

**Research Highlights:** Millimeter-wave radar systems succeeded in the non-contact measurement of respiration in outdoor-housed rhesus macaques from > 5 m. Our results demonstrate the feasibility of radar-based remote monitoring to assess the welfare of zoo-housed animals.

## INTRODUCTION

Vital signs such as hormones, heartbeat, and respiration are pivotal markers for improving animal welfare (Ralph and Tilbrook, 2016). These markers allow for the assessment of mental and physical conditions that may not readily manifest as behaviors. Their utility is extended to various domains, including disease testing, stress evaluation (Heimbürge et al., 2019), and reproductive monitoring (Pickard, 2003). Hormonal analysis based on fecal samples serves as the primary biological indicator for evaluating animal welfare (Hill and Broom, 2009; Binding et al., 2020). Fecal samples are suitable for middle- to long-term monitoring given their capacity to reflect hormone concentrations accumulated over hours to days (Behringer and Deschner, 2017); however, are limited in their ability for the continuous real-time assessment of the mental and physical status of animals.

Respiration is a physiological indicator that can overcome these limitations (Hill and Broom, 2009). Respiration, which is subject to modulation by the autonomic nervous system, exhibits immediate responses to external stimuli, thus facilitating the real-time assessment of the mental and physical conditions of animals (Broom and Johnson, 2019). An illustrative study by Dembiec et al. (2004) demonstrated that the respiratory rate of tigers (*Panthera tigris*) increased significantly during and after transportation between captive environments, suggesting the taxing nature of animal transport. Nevertheless, most vital-sign measurements rely on contact or implantable devices (Aureli et al., 1999; Washer, 2021). The use of such devices results in significant stressors, including anesthesia, capture, or surgery. In addition, the target animals may damage the measurement devices shortly after installation (Hill and Broom, 2009).

Given these limitations, the advancement and application of non-contact vital measurement techniques in captive animals have gained attention. Current approaches include video recordings used to detect subtle alterations in skin color caused by physiological activities (e.g., Unakafov et al., 2018; Zhao et al., 2013; Wang et al., 2023). However, this method has two limitations. First, the necessity of accessing exposed skin for measurements poses challenges in monitoring free-moving animals whose body surfaces are mostly covered with fur or feathers (Al-Naji et al., 2019, as an exception). Second, the susceptibility of skin surface color to ambient light changes complicates the acquisition of reliable data within captive outdoor environments commonly found in zoos.

Respiration measurement using millimeter-wave radar is a promising technique that can overcome these challenges. By emitting radio waves from a radar and receiving the reflected waves from the target, it can successfully measure the respiratory-derived periodic displacements of the skin surface (Koda et al., 2021). This approach has two notable advantages. First, millimeter waves can permeate animal fur to reach the skin surface, enabling measurements in fur-covered sites. The application of millimeter-wave radar for respiration measurements has been successful in some non-human mammals (Matsumoto et al., 2022, 2023; Sakamoto et al., 2023; Tuan et al., 2022). Second, millimeter waves are minimally susceptible to sunlight and other environmental factors (Kurata et al., 2004); therefore, has potential applications in outdoor environments.

However, many previous studies on non-human mammals have involved anesthesia, restraint, or radar systems positioned in close proximity to the targets. In addition, these trials were conducted without direct sunlight exposure. Therefore, it remains unclear whether radar-based non-contact respiratory measurement can be performed in free-moving animals living in outdoor captive enclosures.

In this study, we explored the feasibility of using millimeter-wave radar to remotely measure the respiration of free-moving rhesus macaques (*Macaca mulatta*) in captive outdoor environments from outside their enclosures.

## METHODS

### Study subjects and housing

Captive rhesus macaques (*Macaca mulatta*) from Kyoto City Zoo, Japan were used as the study subjects; 12 individuals were included in the study (11 adult females and one adult male), and only females were used for the measurements. The study population was housed in a circular concrete outdoor enclosure with a diameter of 18 m. Although the area had sufficient sunshade, it was exposed to sunlight on sunny days.

### Data collection

Measurements were conducted on August 2 (Trial 1) and November 17 (Trial 2), 2022, between 9:00 and 11:00. The analyzed measurement durations were 100 s for Trial 1 and 60 s for Trial 2. Table 1 summarizes the weather conditions on the day of each trial. During Trial 1, the target remained in the shaded area, whereas during Trial 2, the target was exposed to sunlight.

**Table 1.** Climatic conditions on measurement dates.

| Date | Weather | Temperature<sup>†</sup> | Relative humidity<sup>†</sup> | Average wind speed<sup>†</sup> |
| --- | --- | --- | --- | --- |
| August 2, 2022 | Sunny | 31.1–33.7 °C | 56–63% | 1.8–2.6 m/s |
| November 17, 2022 | Sunny | 10.5–14.5 °C | 57–74% | 0.6–1.4 m/s |

<sup>†</sup> The range of values between 9:00 and 11:00 obtained from the 10-minute weather data for Kyoto City in the database published by the Japan Meteorological Agency (https://www.data.jma.go.jp/obd/stats/etrn/index.php?prec_no=61&block_no=47759&year=2022&month=&day=&view=).

Radar-based vital measurements are affected by body movement (Spelman & Kindt, 1979). To mitigate this effect, we focused on resting monkeys exhibiting minimal motion. We used a pair of frequency-modulated continuous-wave (FMCW) array radar systems with a center frequency of 79 GHz, center wavelength of 3.8 mm, and bandwidth of 3.6 GHz. Each radar was equipped with three transmitters and four receivers to enhance sensitivity in detecting signals from the target. Radar devices were positioned at the outer edge of the enclosure (Figure 1). The distance between the target and two radar system was 7.28 and 7.68 m for Trial 1 and 5.72 and 5.95 m for Trial 2 (Table 2). Both radar systems emitted radio waves toward the target animal and captured the reflected waves to estimate respiration. Due to ethical considerations, we did not use contact-based respiration measurements to validate the obtained values. Instead, we simultaneously measured the respiration of the same target using two radar systems and examined the Pearson’s correlation coefficients to determine the reliability of our methods.

**Table 2.** Measurement context and results of the two trials.

| Trial | Date | Sunlight on the target body | Posture | Estimated measurement body-site | Distance (m) <sup>†</sup> | Average respiratory rate (bpm) <sup>†</sup> | Correlation coefficient |
| --- | --- | --- | --- | --- | --- | --- | --- |
| 1 | August 2, 2022 | Absent | Lying | Fur-covered right side of the body | 7.28, 7.68 | 55.56, 55.57 | 0.94 |
| 2 | November 17, 2022 | Present | Lying | Fur-covered back | 5.72, 5.95 | 37.75, 39.65 | 0.89 |

<sup>†</sup> Measurements obtained from two radar systems.

**Figure 1.**
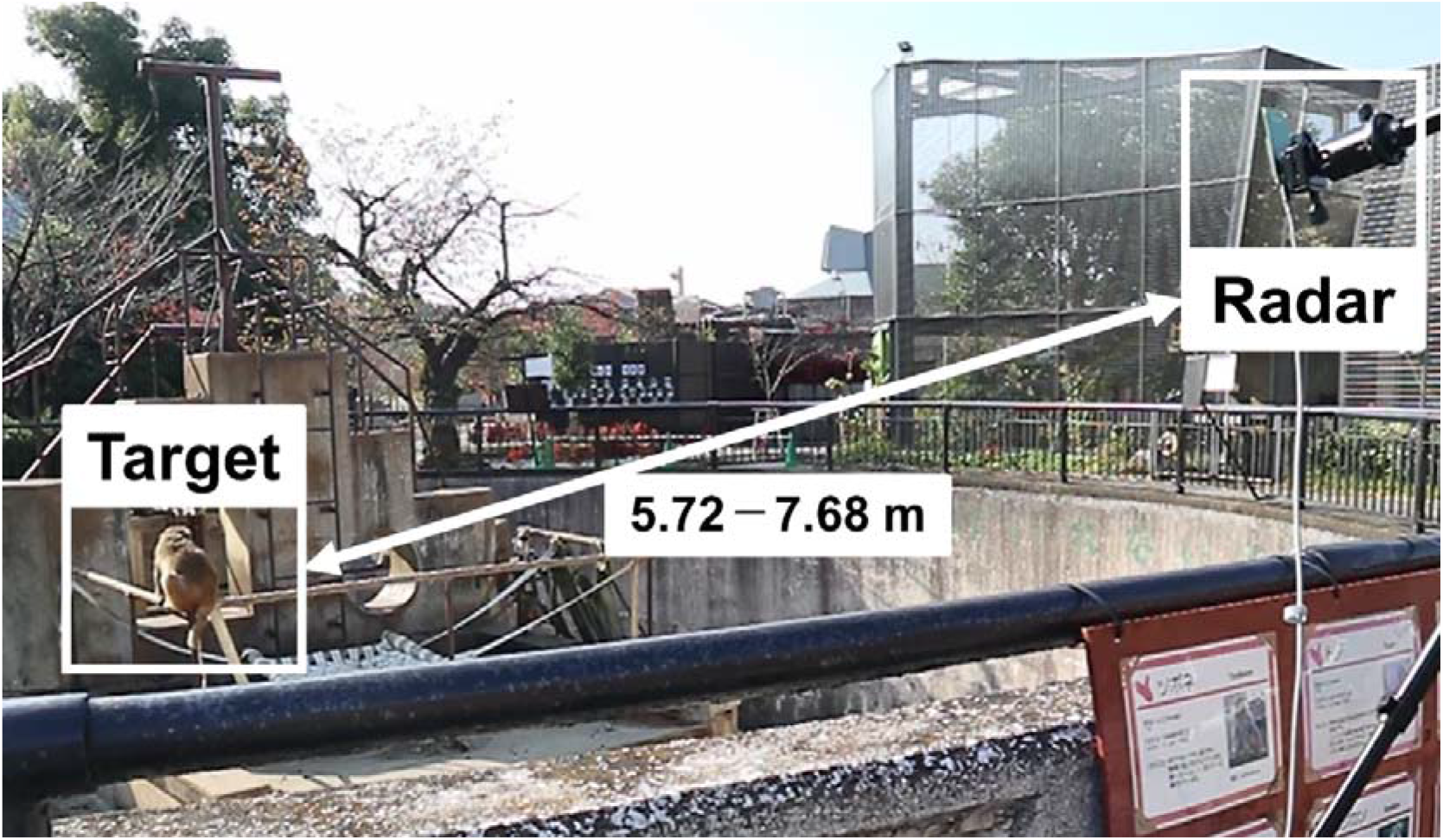
Experimental setup

These settings adhered to the Radio Radiation Protection Guidelines issued by the Ministry of Internal Affairs and Communications, Japan, ensuring minimal impact on the targets. This study was approved by the Ethics Committee of the Kyoto City Zoo (2022-KCZ-006).

### Data analysis

The periodic displacement of the skin surface was extracted from the reflected waves to measure respiration rate and intervals. The first step in the analysis was the construction of a radar image based on reflected waves. This radar image was generated in a Cartesian coordinate framework that encompassed measurement angles and distances. The initial radar images included information on undesired reflected waves (clutter) originating from static objects (such as terrain, barriers, enrichment items) and body movements, along with cyclic displacements attributed to the respiration of the target. To eliminate this clutter, time-averaged components were subtracted from the radar images. Subsequently, the position of the target was estimated by identifying the signal with the highest reflected power in the radar image. Thereafter, *z*(*t*), the displacement waveform of the skin surface, was obtained from the signal phase at the estimated position. Sakamoto et al. (2023) previously described the procedural steps of radar image creation, clutter elimination, and estimation of the skin surface displacement.

During Trial 1, the target exhibited little body movement, allowing for the reduction of undesired components caused by the body motion, using the clutter removal method described above. Therefore, the respiration interval *τ* was determined as the minimal estimates of the evaluation function *F*(*τ*), as follows:

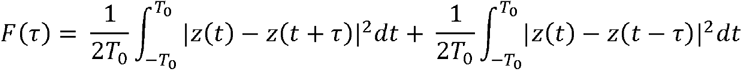

where 2*T*_0_ is the window width (2.2 s in this study), and *t* is the slow time. Since the sampling interval was finite, the point at which *F*(*τ*) exhibited its minimum was not necessarily on the sample point. Therefore, we estimated the parabola passing through *τ* and its front and back, and the point that provided the minimum value in the parabola was estimated as the respiratory interval.

During Trial 2, the target animal was highly mobile, causing undesired components caused by the body motion. Therefore, we used the analyzing method proposed by Sakamoto et al. (2023) to suppress nonperiodic body movement.

## RESULTS

In both trials, we succeeded in observing the periodic displacement of the skin surface (Figure 2). The average respiratory rates estimated from the two radar systems were 55.56 and 55.57 bpm for Trial 1 and 37.75 and 39.65 bpm for Trial 2 (Table 2). The correlation coefficients for the respiratory intervals between the two radar systems were 0.94 (Trial 1) and 0.89 (Trial 2).

**Figure 2.**
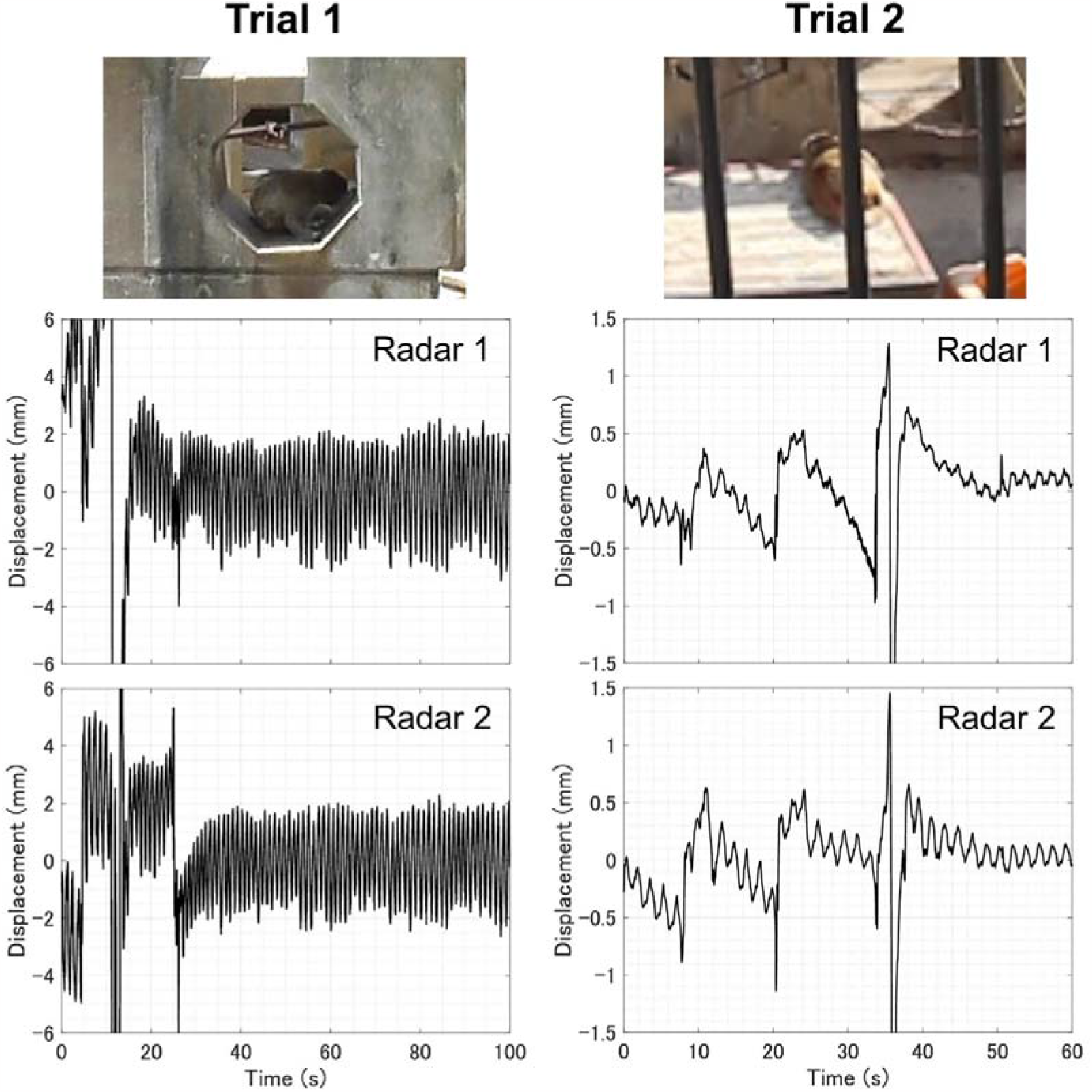
Periodic displacement waveform of the skin surface captured through a pair of millimeter-wave radar systems. The photographs depict the target monkeys during our trials. To enhance the visual clarity of periodic displacements, parts of displacements that exhibited substantial fluctuations due to body movements are not represented on the graph.

In the data analysis, two factors led to data noises in obtaining precise measurements. The first was the significant gross body movements of the targets. Reflected waves induced by bodily motion often relayed information on skin surface movement, masking the skin surface displacement arising from respiration. The second included mobile objects in proximity to the subjects. Information from the reflected waves also included the motion of surrounding objects (e.g., enrichment items), which complicated the detection of respiratory-derived skin surface displacement.

## DISCUSSION

In this study, we succeeded in performing non-contact radar-based respiration measurements of free-moving rhesus macaques housed in a zoo environment, particularly outdoors, from outside the captive area (up to 7.68 m). The recorded mean respiratory rates (Table 2) were consistently within the range reported previously for captive rhesus macaques (35−60 breaths per minute: Brourne, 1975; Fortman et al., 2002; Japanese Society for Laboratory Animal Resources, 2019). Furthermore, the measurements obtained from the two radar systems in Trials 1 and 2 correlated well with each other, despite variations in environmental conditions, such as sunlight (Figure 2; Table 2). These findings indicate the applicability of the millimeter-wave radar for respiration measurements in active rhesus monkeys living outdoors. While still relatively scarce, radar-based vital measurements have been reported in some species (Gong et al., 2022; Matsumoto et al., 2022, 2023; Suzuki et al., 2009; Tuan et al., 2022; Wang et al., 2020; Wang et al., 2022). This suggests that millimeter-wave radar technology for non-contact vital measurements in outdoor settings will pave the way for understanding the physical and mental states of various animal species.

Two factors that may contribute to data noises, reducing the accuracy of measurements, were observed in this study. First, the substantial gross body movements exhibited by the target posed a challenge. Because our technology relies on waves reflected from the skin surface, body movements influence the reflections and result in difficulties in analysis. The measurements in this study were limited to resting individuals. Future research should focus on developing measurement techniques capable of withstanding substantial body movements (Sakamoto et al., 2023).

The second factor was objects in motion other than the subject. The radio waves were not solely reflected by the target, but also by the objects within its vicinity. Although the analytical procedure eliminates reflections from static objects, the removal of reflected waves from objects in considerable motion posed a challenge. Consequently, when moving objects, such as plants or enrichment tools, surround a target, the reflected waves include information relayed from these objects, complicating the detection of skin surface displacements linked to respiration. Furthermore, raindrops may cause distortions in radar reflections, rendering accurate measurements challenging. It is imperative to pay meticulous attention to the captive environment and its conditions when measuring vital signs using radar technology.

Building upon our results, further advancements in non-contact vital measurement technology could significantly contribute to the monitoring of the mental and physical well-being of captive animals to ensure their comfort. For example, Suzuki et al. (2009) installed radar on the ceiling of a zoo enclosure and successfully performed remote and continuous respiration measurements on a hibernating Japanese black bear (*Ursus thibetanus japonicus*) over three months. This enabled caregivers to remotely oversee the health status of hibernating bears, which was characterized by minimal movement, relating to their health. The installation of radar in zoos can facilitate the continuous and real-time assessment of the mental and physical states of animals during their daily lives.

Finally, we believe that our procedures can be applied not only to zoo animals but also to wild animals. Using a radar set at up to 7.68 m from the target, we successfully measured the respiration of free-moving rhesus macaques in an outdoor environment without direct contact. This setup is applicable to wild macaques habituated to human observers. Future studies should conduct measurements on wild macaques living in natural settings to validate the applicability of radar-based non-contact vital measurements in wildlife. Visualizing the physical and mental states of wild animals during their daily lives could not only deepen our understanding of the species but also significantly enhance animal welfare in captive settings.

## Acknowledgments

We are grateful to the staff of Kyoto City Zoo for their invaluable assistance in facilitating this study. We thank Dr. Hirofumi Taki and Dr. Shigeaki Okumura of MaRI Co., Ltd. for their expert guidance. This study was financially supported by the SECOM Science and Technology Foundation (to TS); JSPS KAKENHI under Grants 19H02155, 21H03427, and 23H01420 (to TS); JST under Grant JPMJMI22J2 (to TS and MM); and JST SPRING JPMJST2110 (to TM).

